# STELAR: A statistically consistent coalescent-based species tree estimation method by maximizing triplet consistency

**DOI:** 10.1101/594911

**Authors:** Mazharul Islam, Kowshika Sarker, Trisha Das, Rezwana Reaz, Md. Shamsuzzoha Bayzid

**Affiliations:** Department of Computer Science and Engineering, Bangladesh University of Engineering and Technology, 1205 Dhaka, Bangladesh; Department of Computer Science, The University of Texas at Austin, 78712 Texas, USA

**Keywords:** phylogenomics, multi-species coalescent process, gene tree incongruence, incomplete lineage sorting

## Abstract

**Background:** Species tree estimation is frequently based on phylogenomic approaches that use multiple genes from throughout the genome. However, estimating a species tree from a collection of gene trees can be complicated due to the presence of gene tree incongruence resulting from incomplete lineage sorting (ILS), which is modelled by the multi-species coalescent process. Maximum likelihood and Bayesian MCMC methods can potentially result in accurate trees, but they do not scale well to large datasets.

**Results:** We present STELAR (Species Tree Estimation by maximizing tripLet AgReement), a new fast and highly accurate statistically consistent coalescent-based method for estimating species trees from a collection of gene trees. We formalized the constrained triplet consensus (CTC) problem and showed that the solution to the CTC problem is a statistically consistent estimate of the species tree under the multi-species coalescent (MSC) model. STELAR is an efficient dynamic programming based solution to the CTC problem which is highly accurate and scalable. We evaluated the accuracy of STELAR in comparison with SuperTriplets, which is an alternate fast and highly accurate triplet-based supertree method, and with MP-EST and ASTRAL – two of the most popular and accurate coalescent-based methods. Experimental results suggest that STELAR matches the accuracy of ASTRAL and improves on MP-EST and SuperTriplets.

**Conclusions:** Theoretical and empirical results (on both simulated and real biological datasets) suggest that STELAR is a valuable technique for species tree estimation from gene tree distributions.

## Background

Estimated species trees are useful in many biological analyses, but accurate estimation of species trees can be quite complicated. Species tree inference can potentially result in accurate evolutionary history using data from multiple loci. Therefore, due to the advent of modern sequencing technologies, it is increasingly common to infer trees by analyzing sequences from multiple loci. However, combining multi-locus data is difficult, especially in the presence of gene tree discordance [1]. A traditional approach to species tree estimation is called concatenation (also known as combined analysis), which concatenates gene sequence alignments into a supergene matrix, and then estimates the species tree using a sequence based tree estimation technique (e.g., maximum parsimony, maximum likelihood etc.). Although it has been used in many biological analyses, concatenation can be problematic as it is agnostic to the topological differences among the gene trees, can be statistically inconsistent [2], and can return incorrect trees with high confidence [3, 4, 5, 6].

Recent modeling and computational advances have produced methods that explicitly take the gene tree discordance into account while combining multi-locus data to estimate species trees. Incomplete lineage sorting (ILS) (also known as deep coalescence) is one of the most prevalent reasons for gene tree incongruence [1], which is modelled by the MSC [7]. Rapid radiation is known to happen in many organisms (for example, birds) where ILS is likely to occur [8]. Due to this growing awareness that ILS can be present in many phylogenomic datasets, ILS-aware species tree estimation techniques have gained substantial attention from systematists. These types of method are usually known as “summary methods” as they summarize a set of gene trees to estimate the species trees [9].

Several species-tree estimation methods have been proven to be statistically consistent under the multi-species coalescent model, meaning that they will return the true species tree with high probability given a sufficiently large number of true gene trees sampled from the distribution defined by the species tree. Methods that are statistically consistent under the MSC include ASTRAL [10], MP-EST [11], *BEAST [12], NJst [13], BUCKy [14], GLASS [15], STEM [16], SNAPP [17], SVDquartets [18], STEAC [19] and ASTRID [20]. *BEAST, which is a Bayesian technique, can co-estimate both gene trees and species trees, and simulation studies suggest that *BEAST can be considered the best of the co-estimation methods with excellent accuracy, but it is computationally very intensive to run on even moderately-sized dataset [21, 9]. SuperTriplets [22] is a triplet-based supertree method which tries to find an asymmetric median supertree according to a triplet dissimilarity. SuperTriplets is very fast and accurate and was used in a number of important phylogenomic studies [23, 24, 25, 26, 27, 28]. ASTRAL and MP-EST are two of the most accurate and widely used summary methods that are much faster than *BEAST. ASTRAL has been shown to be faster and more accurate than MP-EST and can handle large dataset containing hundreds of species which is not possible by MP-EST [10, 29].

ASTRAL finds a species tree that maximizes the number of consistent quartets between the species tree and the gene trees. MP-EST maximizes a pseudo-likelihood estimate utilizing the underlying triplet distribution of the gene trees. Quartet and triplet based methods are robust to the “anomaly zone” [30, 31] (a condition where the most probable gene tree topology may not be identical to the species tree topology) as there are no anomalous rooted three-taxon species trees [30, 32] and no anomalous unrooted four-taxon species trees [33, 31].

We present STELAR (Species Tree Estimation by maximizing tripLet AgReement), a new coalescent-based method which finds a species tree that agrees with the largest number of triplets induced by the gene trees. STELAR is statistically consistent under the MSC model, fast (having a polynomial running time), and highly accurate – enabling genome wide phylogenomic analyses. We report, on an extensive experimental study, the performance of STELAR in comparison with ASTRAL, MP-EST and SuperTriplets. These quartet- and triplet-based methods are known to be highly accurate and reliable, and are being widely used. Our experiments showed that STELAR is as good as ASTRAL, and is better than MP-EST and SuperTriplets in most of the model conditions. Thus, with the desirable property of being statistically consistent and reliable performance on challenging realistic model conditions, STELAR will be a useful tool for genome-scale analyses of species phylogeny.

## Methods

Design of ASTRAL and other quartet-based methods [10, 34] and their proofs of being statistically consistent are based upon the fact that unrooted 4-taxon species trees do not contain anomaly zone [33, 31]. We use similar design in STELAR, utilizing the fact that rooted 3-taxon species trees do not contain anomaly zone [30, 32].

### Definitions and notation

Let *T* be a binary rooted tree, leaf-labelled by species set 𝓧 with *n* taxa. Each internal node *u* in a tree *T* divides the set of taxa present in the subtree rooted at *u* into two distinct sets. We call this a *subtree-bipartition*, denoted by *SBP_T_* (*u*), which was originally defined in [35]. We denote by *T*_*u*_ the subtree under node *u* of tree *T*. We denote the leaves in *T* by *L*(*T*), an arbitrary set of three species {*a, b, c*} ⊂ 𝓧 by *r*, and a rooted topology on *r* by *t*_*r*_. We use *a|bc* to show that *b* and *c* are siblings. Possible three topologies on {*a, b, c*} are: *a|bc, b|ca, c|ab*. The probability, under the MSC model, of observing an induced gene tree triplet that matches the species tree topology is higher, and the other two triplet topologies are less probable and equal to each other [30]. The triplet tree topology that appears more frequently than the two alternative topologies is called the *dominant* topology. 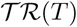 denotes the set of 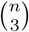 triplet topologies induced by the tree *T*. We denote by *T |r* the triplet tree topology obtained by restricting *T* to the three species in *r*.

### Problem definition

A natural optimization problem for estimating a species tree from a collection of gene trees under triplet consistency would be first decomposing the input gene trees into their induced triplets, and then estimating a species tree by amalgamating the dominant triplets [36]. Building such a consensus tree from a set of triplets has been shown to be an NP-hard problem, and exact and heuristic algorithms have been proposed [37, 38]. This approach considers only the dominant triplets which could be problematic when the frequencies of the alternate triplet topologies are very close to that of the dominant one, and when all the dominant triplets are not compatible with each other meaning that all of them do not agree with a single species tree. Moreover, this approach has to explicitly enumerate and consider 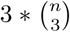 possible triplets, making it computationally expensive for larger values of *n*. An alternate approach would be to consider the relative frequencies of all the triplets, and infer a species tree that agrees with the largest number of triplets induced by the gene trees. Methods that do not explicitly decompose the gene trees into induced triplets and quartets are preferred over the ones that take a set of quartets or triplets as input, since the latter types of methods demand high computational time and memory, and thus are not scalable to large dataset [34, 10]. This is possibly one of the reasons why quartet amalgamation method like QFM [34], despite having excellent accuracy, is not as popular as ASTRAL. Taking these issues into consideration, we introduce a constrained version of the triplet consistency problem, which we call the *Constrained Triplet Consensus* (CTC) problem, and provide efficient exact and heuristic solutions to the CTC without having to decompose the gene trees into their induced triplets. We formalize the CTC problem as follows.

- **Input**: a set 𝓖 of rooted gene trees on 𝓧 and a set 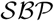 of subtree-bipartitions.
- **Output**: a species tree *ST* on 𝓧 that maximizes the triplet agreement with 𝓖 and all its subtree-bipartitions are in 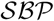.

Note that when 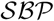 is set to all possible subtree-bipartitions on 𝓧, the triplet consistency problem will find the globally optimal solution (exact solution). Otherwise, when 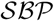 contains only the subtree-bipartitions in the input gene trees, the search space will include only those species trees where every subtree-bipartition is present in at least one of the gene trees in 𝓖, and we call this the constrained version or heuristic version.

**Theorem 0.1** *Given a set 𝓖 of true gene trees, solution to the CTC problem (both exact and constrained version) is a statistically consistent estimator of the species tree topology under the MSC model*.

Proof Let 𝓖 be a set of sufficiently large number of true gene trees that has evolved within the model species tree *T* under the multi-species coalescent model. We know that a rooted 3-taxon species tree does not have any anomaly zone [32]. So, as we increase the number of gene trees, each triplet topology induced by the species tree will have more frequency in 𝓖 than its alternatives, with probability that approaches 1. Let *w_𝓖_* (*t*_*r*_) be the number of trees in 𝓖 that agree with *t*_*r*_. Hence, for every triplet *r* and every possible tree *T*′, *w_𝓖_* (*T|r*) *≥ w_𝓖_* (*T*′|*r*) with probability that approaches 1 for a large enough number of gene trees. Therefore, if 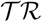 is the set of all possible triplets on 𝓧, then

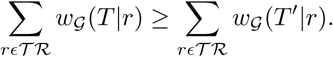

By definition, the exact solution to the CTC problem will find a tree that will maximize the triplet agreement with 𝓖. Therefore, for a sufficiently large number of true gene trees, the optimal tree under CTC will be identical to the true species tree *T* with probability approaching one.

We now show that the solution to the constrained version is also statistically consistent. If we increase the number of gene trees, with probability approaching 1, at least one of the gene tree topologies in 𝓖 will be identical to the the true species tree topology. Therefore, with probability that tends to 1, the set 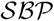 will contain all the subtree bipartitions from the true species tree, and therefore the solution to the CTC problem will be identical to the true species tree *T*.

### Algorithmic design of STELAR

We propose a dynamic programming (DP) solution to the CTC problem. The algorithmic design (and corresponding theoretical results) in STELAR is structurally similar to ASTRAL. This sort of DP-based approach, which implicitly finds a maximum or minimum clique in a graph modelled from the input gene trees, was first used by [39] and later was used in Phylonet [40, 41], DynaDup [35, 42] and ASTRAL. The key idea in STELAR is to find an equation for computing the number of triplets in the gene trees that agree with a given subtree in the species tree, which ultimately enables us to design a dynamic programming algorithm for estimating a species tree by maximizing the triplet agreement.

We call a triplet *t*_*r*_ = *a|bc* to be mapped to a subtree-bipartition at an internal node *x* in 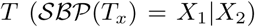 when {*a, b, c*} ⊆ *L*(*T*_*x*_), and the topology of *t*_*r*_ is compatible (consistent) with *T*_*x*_ (see Fig. 1). Each triplet mapped to a subtree-bipartition (*X*_1_*|X*_2_) will have either two leaves from *X*_1_ and one leaf from *X*_2_ or two leaves from *X*_2_ and one leaf from *X*_1_. Therefore, we have the following results.

**Figure 1.**
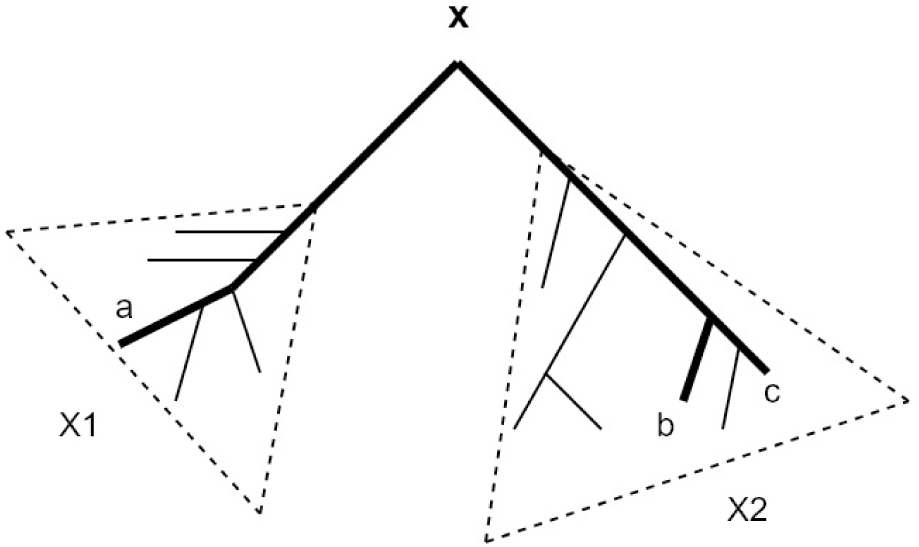
Mapping of a triplet to a subtree-bipartition. Each internal node *x* in a rooted tree defines a subtree-bipartition (X_1_|X_2_). Each induced triplet in *T* maps to a subtree-bipartition in *T*. This figure shows the mapping of a triplet a|bc to a subtree-bipartition *x* = X_1_|X_2_.

**Lemma 0.2** *Total number of triplets mapped to a subtree-bipartition x* = (*X*_1_*|X*_2_) *of a rooted binary tree T, where |X*_1_*|* = *n*_1_ *and |X*_2_*|* = *n*_2_ *is*:

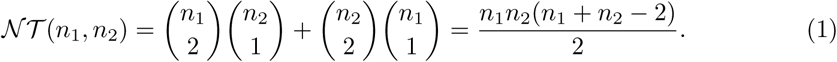

**Lemma 0.3** *For two subtree-bipartitions x* = (*X*_1_*|X*_2_) *and y* = (*Y*_1_*|Y*_2_) *where* (*X*_1_ ∪ *X*_2_) *⊆ 𝓧 and* (*Y*_1_ ∪ *Y*_2_) *⊆ 𝓧, the number of triplets mapped to both x and y is given by*:

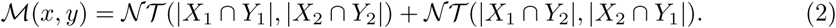

*Proof* We can obtain two possible subtree bipartitions by taking pairwise intersection: *z*_1_ = (*X*_1_ ∩ *Y*_1_*|X*_2_ ∩ *Y*_2_) and *z*_2_ = (*X*_1_ ∩ *Y*_2_*|X*_2_ ∩ *Y*_1_). Note that each leaf in *z*_*i*_ (*i ∈* 1, 2) is also a leaf in both *x* and *y*, and so each triplet mapped to *z*_*i*_ is also mapped to *x* and *y* and the number of mapped triplets can be computed by Eqn. 1.

We now prove that Eqn. 2 counts a mapped triplet only once. Without loss of generality, let *t*_*r*_ = *ab|c* be mapped to both *x* and *y* where *a, b* ∈ *X*_1_, *c* ∈ *X*_2_, *a, b* ∈ *Y*_1_ and *c* ∈ *Y*_2_. Eqn. 2 will count *t*_*r*_ for *z*_1_ = (*X*_1_ ∩ *Y*_1_*|X*_2_ ∩ *Y*_2_) only as *z*_2_ will have no element from {*a, b, c*}. Thus the lemma follows.

We now see how to compute the triplet consistency (*TC*) score of a species tree *ST* with respect to a set 𝓖 of gene trees, which we denote by *TC_𝓖_* (*ST*). If 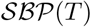 is the set of subtree-bipartitions in *T*, for a subtree-bipartition *x* and a set of rooted binary trees 𝓖, the total number of triplets in 𝓖 that are mapped to *x* is given by:

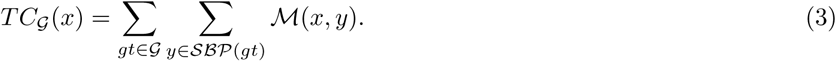

Thus, the *TC* score of an estimated species tree *ST* is obtained by,

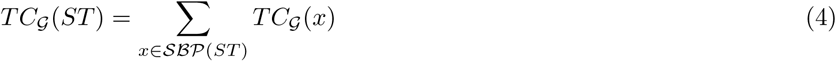

Therefore, we can design a dynamic programming approach which starts with the whole species set 𝓧 and recursively divides it into smaller subsets and picks the partition that maximizes the triplet consistency. At each level, we compute *V* (*A*) which gives the score of an optimal subtree on leaf set *A* ⊆ 𝓧. At the end level, we have computed *V* (𝓧) and sufficient information so that backtracking yields the corresponding subtree-bipartitions in the optimal species tree.

### Dynamic Programming

#### Base case

*|A|* = 1; *V* (*A*) = 0

#### Recursive relation

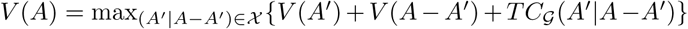, where 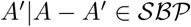.

### Running time analysis

**Lemma 0.4** *Given a set 𝓖 of k gene trees on n taxa and a subtree-bipartition x, computing the triplet consistency score TC_𝓖_* (*x*) *takes O*(*n*^2^*k*) *time*.

Proof A gene tree with *n* taxa contains *O*(*n*) subtree-bipartitions. Thus, for a set of *k* input gene trees, there will be *O*(*nk*) subtree-bipartitions. To find the triplet consistency score *TC*_𝓖_(*x*) of a subtree-bipartition *x* using Eqn. 3, we need to score 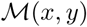 for all subtree-bipartions *y* present in the input gene trees. Since we represent each subtree-bipartion by a bitset, calculating 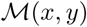 (using Eqn. 2) takes *O*(*n*) time. Since there are *O*(*nk*) subtree-bipartions in 𝓖, calculating *TC*_𝓖_ (*x*) takes *O*(*n*^2^*k*) time.

**Theorem 0.5** *For a given set 𝓖 of k gene trees on n taxa and a set 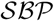 of subtree-bipartitions, STELAR runs in* 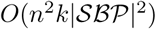 *time*.

Proof For a specific cluster *A*, *V* (*A*) can be computed in 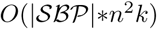 time since at worst we need to look at every subtree-bipartition in 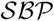, and computing *TC*_𝓖_ (*x*) takes *O*(*n*^2^*k*) time. Note that this is a conservative analysis since the number of subtree-bipartitions that the DP algorithm needs to consider for computing *V* (*A*) for a specific cluster *A* is, in reality, much less than 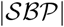 (especially for *A* where *|A| < |𝓧|*). There will be 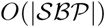 possible clusters for which we need to compute *V* (*A*). Thus, the running time of STELAR is 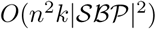

## Results

### Experimental Studies

We compared STELAR with SuperTriplets, which is a very fast and highly accurate triplet based supertree technique and has been used in several important phylogenomic studies [22, 23, 24, 25, 26, 27]. We also evaluated STELAR in comparison with ASTRAL and MP-EST, two of the most widely used and accurate coalescent-based methods. ASTRAL has already been compared and shown to outperform many of the existing species tree estimation methods (e.g., BUCKy-pop, MRP, greedy consensus, CA-ML, etc.), especially in the presence of moderate to high levels of ILS. So we did not include those methods in this study. Most of the datasets analyzed in this study are prohibitively large for *BEAST to analyze. We ran MP-EST with 10 random starting points and selected the species tree with the highest pseudo-likelihood value.

We used previously studied simulated and biological datasets to evaluate the performance of these methods. We compared the estimated species trees to the model species tree (for the simulated datasets) or to the scientific literature (for the biological datasets), to evaluate accuracy. We used normalized Robinson-Foulds distance (RF rate) [43] to measure the tree error. The RF distance between two trees is the sum of the bipartitions (splits) induced by one tree but not by the other, and vice versa. All the trees estimated by ASTRAL, STELAR and MP-EST in this study are binary and so RF rate, False Positive (FP) rate and False Negative (FN) rate are identical. However, the trees estimated by SuperTriplets may produce multifurcations (unresolved trees). We performed Wilcoxon signed-rank test (with *α* = 0.05) to measure the statistical significance of the differences between two methods.

### Datasets

We studied three collections of simulated datasets: one based on a biological dataset (37-taxon mammalian dataset) that was generated and subsequently analyzed in prior studies [10, 44, 29], and two other simulated datasets from [45] and [44]. Table 1 presents a summary of the datasets analyzed in this paper, showing various model conditions with varying numbers of taxa (11 to 37), ILS levels (reflected in the average topological distance between true gene trees and true species tree) and gene sequence lengths. All datasets consist of gene sequence alignments generated under a multi-stage simulation process that begins with a species tree, simulates gene trees down the species tree under the multi-species coalescent model (and so can differ topologically from the species tree), and then simulates gene sequence alignments down the gene trees.

**Table 1.** Properties of the simulated datasets. Level of ILS is presented in terms of the average topological distance between true gene trees and true species tree.

| Dataset | ILS level | # genes | # sites | # Replicates | Ref. |
| --- | --- | --- | --- | --- | --- |
| 11-taxon | 85% | 5 - 100 | 2000 | 20 | [45] |
| 15-taxon | 82% | 100 - 1000 | 100 - 1000 | 10 | [44] |
| 37-taxon | 18%, 32%, 54% | 25 - 800 | 250 -1000 | 20 | [44] |

Mammalian dataset was simulated by taking the species tree estimated by MPEST on the biological dataset studied in [46]. This species tree had branch lengths in coalescent units, which we used to produce a set of gene trees under the coalescent model. Thus, the mammalian simulation model tree has an ILS level based on a coalescent analysis of the biological mammalian dataset, and other properties of the simulation that are set to reflect the biological sequences [46] studied. We explored the impact of varying amounts of ILS, varying numbers of genes (25 *∼* 800), the impact of phylogenetic signal by varying the sequence length (250, 500, and 1000 base pair (bp)) for the markers. Three levels of ILS were simulated. The basic model species tree has branch lengths in coalescent units, and we produced other model species trees by multiplying or dividing all internal branch lengths in the model species tree by two. This rescaling varies the amount of ILS (shorter branches have more ILS), and also impacts the amount of gene tree estimation error in the estimated gene trees. The basic model condition with moderate amount of ILS is referred to as 1X and the model conditions with higher and lower amounts of ILS are denoted by 0.5X and 2X, respectively.

We used two other simulated datasets: 11-taxon dataset (generated by [45] and subsequently studied by [9, 47]), and 15-taxon datasets (generated and studied by [44]). 11-taxon dataset was simulated under a complex process to ensure substantial heterogeneity between genes and to deviate from the molecular clock [45]. We analyzed the model condition with high amount of ILS as the model condition with lower amount of ILS is very easy to analyze and even methods without any statistical guarantee produced highly accurate tree with just 25 *∼* 50 gene trees [9]. 15-taxon datasets vary in sequence lengths and has high amount of ILS.

We used two biological datasets: the 37-taxon mammalian datasets studied by [46] with 424 genes, and the Amniota dataset from [48] containing 16 species and 248 genes.

### Results on mammalian simulated dataset

We analyzed the performance of SuperTriplets, ASTRAL, MP-EST and STELAR on various model conditions with varying amounts of ILS, numbers of genes and lengths of the sequences. Figure 2(a) shows the average RF rates of three methods on three model conditions with varying amounts of ILS. For all methods, RF rates increase as we increase the amount of ILS. SuperTriplets produced significantly less accurate trees than the other methods, and the trees are not fully resolved. For the dataset with varying amounts of ILS, SuperTriplets produced trees with RF rates 10% *∼* 18%, whereas the error rates of STELAR, MP-EST and ASTRAL range from 4% *∼* 6%. STELAR and ASTRAL are better than MP-EST on all amounts of ILS and, in some cases, the differences are statistically significant (*P <* 0.05). STELAR and ASTRAL are comparable in terms of tree accuracy and no statistically significant difference was noticed.

**Figure 2.**
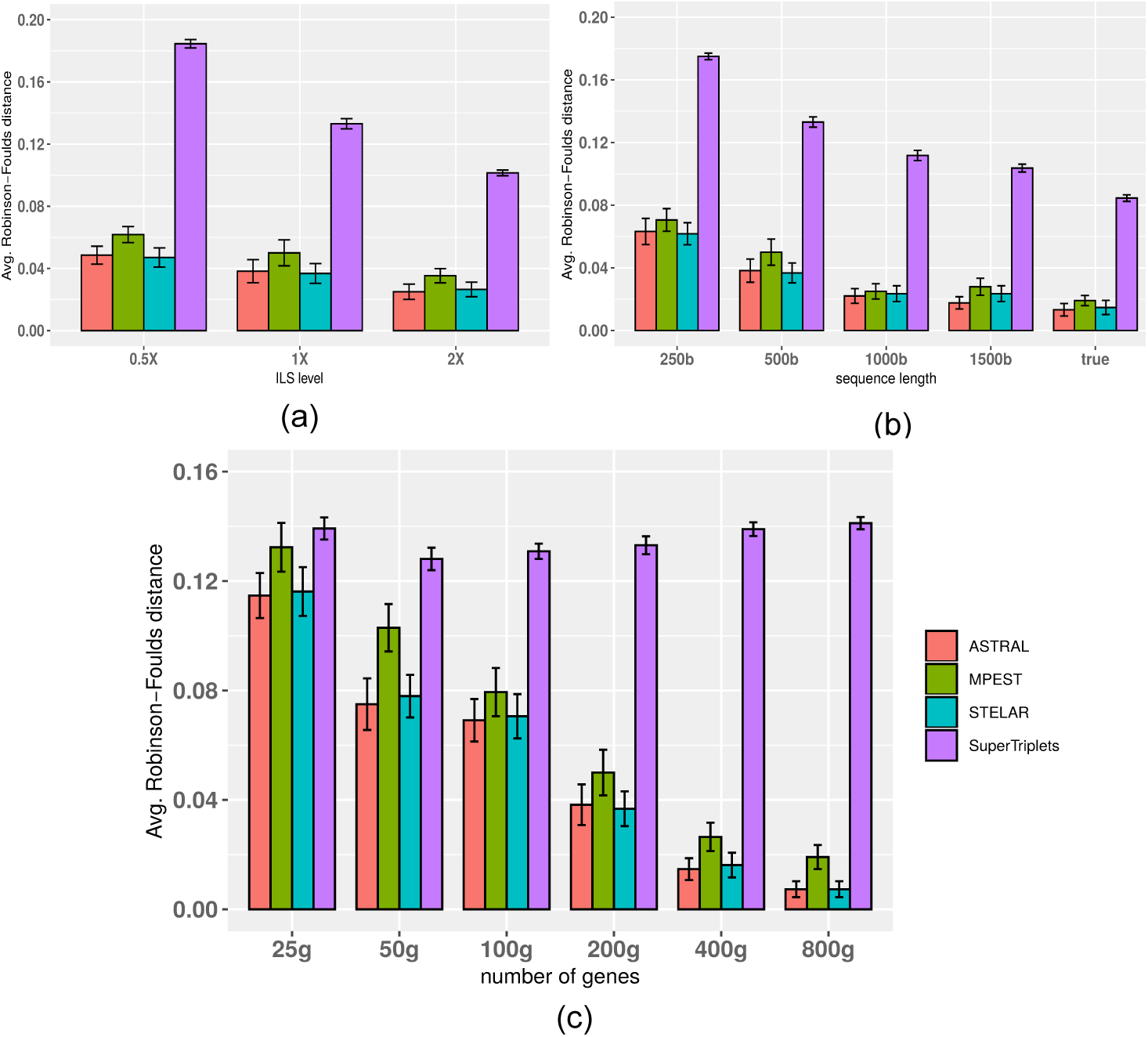
Comparison of ASTRAL, MP-EST, STELAR and SuperTriplets on 37-taxon simulated mammalian dataset. We show the average RF rates with standard error bars over 20 replicates. (a) We fixed the sequence length to 500 bp and number of genes to 200, and varied the amounts of ILS. 2X model condition contains the lowest amount of ILS while 0.5X refers to the model condition with the highest amount of ILS. (b) We varied the amount of gene tree estimation error by varying the sequence lengths from 250 to 1000 bp, while keeping the ILS level (moderate) and the number of genes (200) fixed. (c) We fixed the sequence length to 500 bp and amount of ILS to 1X, and varied the numbers of genes (25, 50, 100, 200, 400, 800).

Figure 2(c) shows the impact of the numbers of gene trees on the accuracy of the estimated species trees. STELAR, MP-EST and ASTRAL improved with increasing numbers of genes (this is expected as these methods are statistically significant and thus increasing the numbers of genes has resulted into improved topological accuracy). However, SuperTriplets failed to leverage the increasing amount of data since it did not improve as we increased the numbers of genes. Moreover, even with 800 genes, SuperTriplets produced unresolved trees. STELAR and ASTRAL achieved similar accuracy and performed significantly better than MP-EST and SuperTriplets across all model conditions.

### Results on 11-taxon dataset

We analyzed both the estimated and true gene trees. In both cases, we varied the number of genes from (5 *∼* 100). Figure 3 shows the results on 11-taxon dataset for both estimated and true gene trees. Similar to the mammalian dataset, SuperTriplets produced the least accurate trees. As expected, all methods improved with increased numbers of genes (except for a few cases for SuperTriplets), and had their best accuracies on true gene trees. The accuracies of ASTRAL, MP-EST and STELAR are almost similar with no statistically significant difference. On true gene trees, ASTRAL, MP-EST and STELAR recovered the true species trees across all the replicates with 50 or higher numbers of genes. However, the accuracy of SuperTriplets did not achieve any notable improvement as we increased the number of genes beyond 25. Unlike mammalian dataset, MP-EST was slightly better than ASTRAL and STELAR on a couple of model conditions (albeit the differences are not statistically significant with *P*-values greater than 0.05).

**Figure 3.**
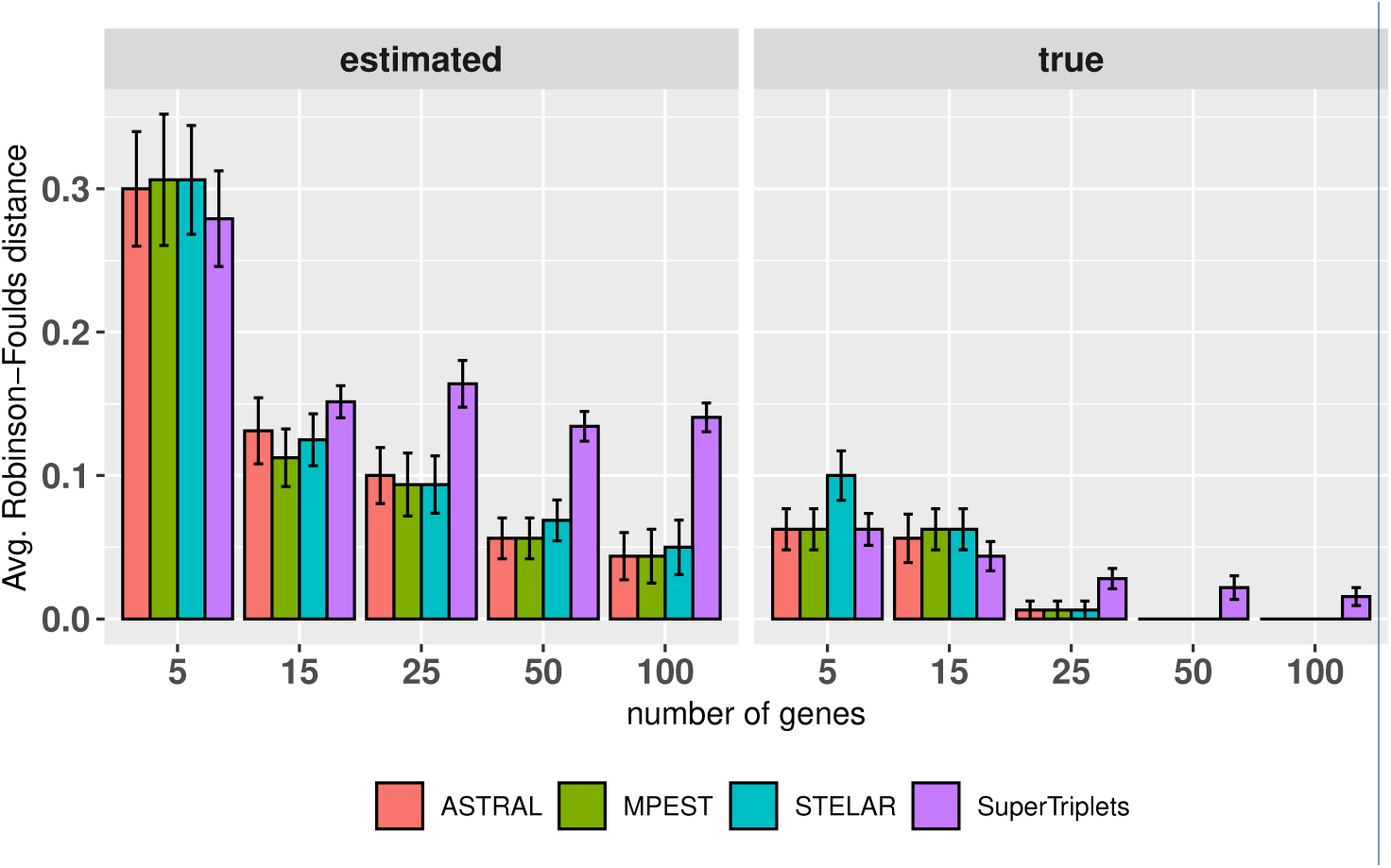
Average RF rates of ASTRAL, MP-EST, STELAR and SuperTriplets on the 11-taxon dataset with high amount of ILS. We varied the numbers of genes from 5 to 100. We analyzed both the estimated and true gene trees. We show the average RF rates with standard error bars over 20 replicates.

### Results on 15-taxon dataset

15-taxon dataset evolved under a caterpillar model species tree with very high level of ILS [44]. We explored the performance on varying sequence lengths (100 bp and 1000 bp), and numbers of genes (100 and 1000). Unline other two dataset, SuperTriplets returned well resolved trees without multi-furcations and matched the accuracy of other methods on this particular dataset. All the methods improved as the the number of genes and sequence length increased, and had their best accuracy (RF rate = 0) on true gene trees (Fig. 4). No statistically significant difference was observed between STELAR and ASTRAL. MP-EST consistently had the highest error rates across all the model conditions on estimated gene trees. On true gene trees, all the methods returned the true species tree on almost all the replicates.

**Figure 4.**
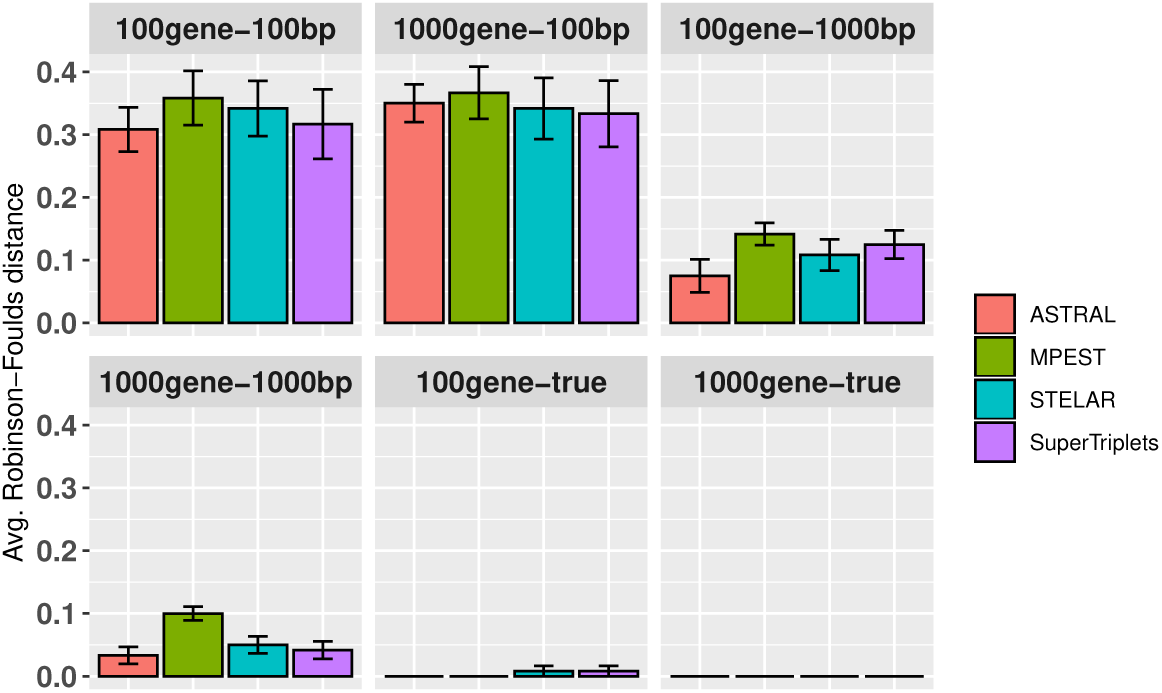
Comparison of ASTRAL, MP-EST, STELAR and SuperTriplets on the 15-taxon dataset. We varied the numbers of estimated gene trees (100 and 1000) and sequence lengths (100 bp and 1000 bp). We also analyzed the true gene trees. We show the average RF rates with standard errors over 10 replicates.

### Results on Biological datasets

#### Amniota dataset

We analyzed the Amniota dataset from [48] containing 248 genes across 16 amniota taxa in order to resolve the position of turtles within the amniotes, especially relative to birds and crocodiles. We re-analyzed both the amino acid (AA) and DNA gene trees, obtained from the authors, using ASTRAL, MPEST, STELAR and SuperTriplets. The sister relationship of crocodiles and birds (forming Archosaurs), and placement of the turtles as the sister clade to Archosaurs are considered reliable and were supported by previous studies [49, 50, 51].

For amino acid dataset, all the methods placed birds and crocodiles as sister groups (forming Archosaurs) and turtles as sister to Archosaurs clade. The unrooted version of STELAR estimated tree is identical to ASTRAL. MP-EST also returned a highly similar tree except for different resolutions within the turtles. SuperTriplets produced an unresolved tree with several multifurcations, leaving bird/turtle/crocodile relationship uncertain. It also did not resolve the Squamates clade.

STELAR and MP-EST on the DNA gene trees produced the same tree which resolved bird/turtle/crocodile relationship as (birds,(turtles,crocodiles)) and thus did not form Archosaurs. However, ASTRAL on the DNA gene trees resolved the relationships as (birds,(turtles,crocodiles)). Therefore, maximizing triplet agreement and quartet agreement both recovered the (birds,(turtles,crocodiles)) relationships on AA gene trees, but differed on the DNA gene trees. Similar to the AA dataset, SuperTriplets did not resolve the bird/turtle/crocodile relationship. We estimated the quartet scores and the triplet scores of these different tree topologies with respect to the corresponding gene trees (see Tables 2 and 3). As ASTRAL estimated trees need to be interpreted as unrooted trees, we root the ASTRAL trees using the correct outgroup (*Protopterus annectens*) in order to compute the triplet scores. As expected, ASTRAL achieved the highest quartet agreement scores and STELAR achieved the highest triplet agreement scores.

**Table 2.**
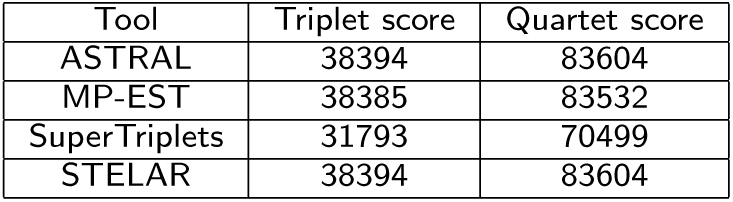
Triplet and quartet scores of the species trees estimated by ASTRAL, MP-EST, SuperTriplets and STELAR on the amino acid (AA) gene trees in Amniota dataset.

| Tool | Triplet score | Quartet score |
| --- | --- | --- |
| ASTRAL | 38394 | 83604 |
| MP-EST | 38385 | 83532 |
| SuperTriplets | 31793 | 70499 |
| STELAR | 38394 | 83604 |

**Table 3.** Triplet and quartet scores of the species trees estimated by ASTRAL, MP-EST, SuperTriplets and STELAR on the DNA gene trees in Amniota dataset.

| Tool | Triplet score | Quartet score |
| --- | --- | --- |
| ASTRAL | 42419 | 97890 |
| MP-EST | 42448 | 97744 |
| SuperTriplets | 40095 | 94121 |
| STELAR | 42448 | 97744 |

#### Mammalian dataset

We re-analyzed the mammalian dataset from [46] containing 447 genes across 37 mammals after removing 21 mislabeled genes (confirmed by the authors), and two other outlier genes. The placement of tree shrews (*Tupaia belangeri*) is of great importance here as previous studies found that the phylogenetic placement of tree shrews is unstable as they can be classified as a sister group to Primates or to Glires [46, 52, 53, 54]. Our analyses using ASTRAL, MP-EST and STELAR recovered the same topology which placed tree shrews as sister to Glires (see Fig. 6). This is consistent to the tree estimated by CA-ML (reported in [46]). However, previous studies with MP-EST, using multi-locus bootstrapping [55], on the same dataset recovered a tree which placed tree shrews as the sister to Primates [54, 10]. SuperTriplets produced a highly similar tree except that it resulted into an unresolved tree with several multi-furcations. It recovered the major clades such as Primates, Glires, Cetartiodactyla, Eulipotyphla, Chiroptera. However, SuperTriplets left the relationships within Primates/Glires/Scandentia unresolved. This is in line with the tree estimated by SuperTriplets on the OrthoMaM database [56], which was reported in [22]. We investigated the quartet and triplet scores of the two alternative hypotheses (regarding the placement of tree shrew) and found that the tree that suggests (tree shrew, Glires) clade has higher quartet and triplet scores (25526915 and 3042547, respectively) compared to the one that suggests (tree shrew, Primates) relationship which yields 25511503 and 3041747 quartet and triplet scores respectively. Therefore, our analyses strongly support the placement of tree shrews as sister to Glires.

**Figure 5.**
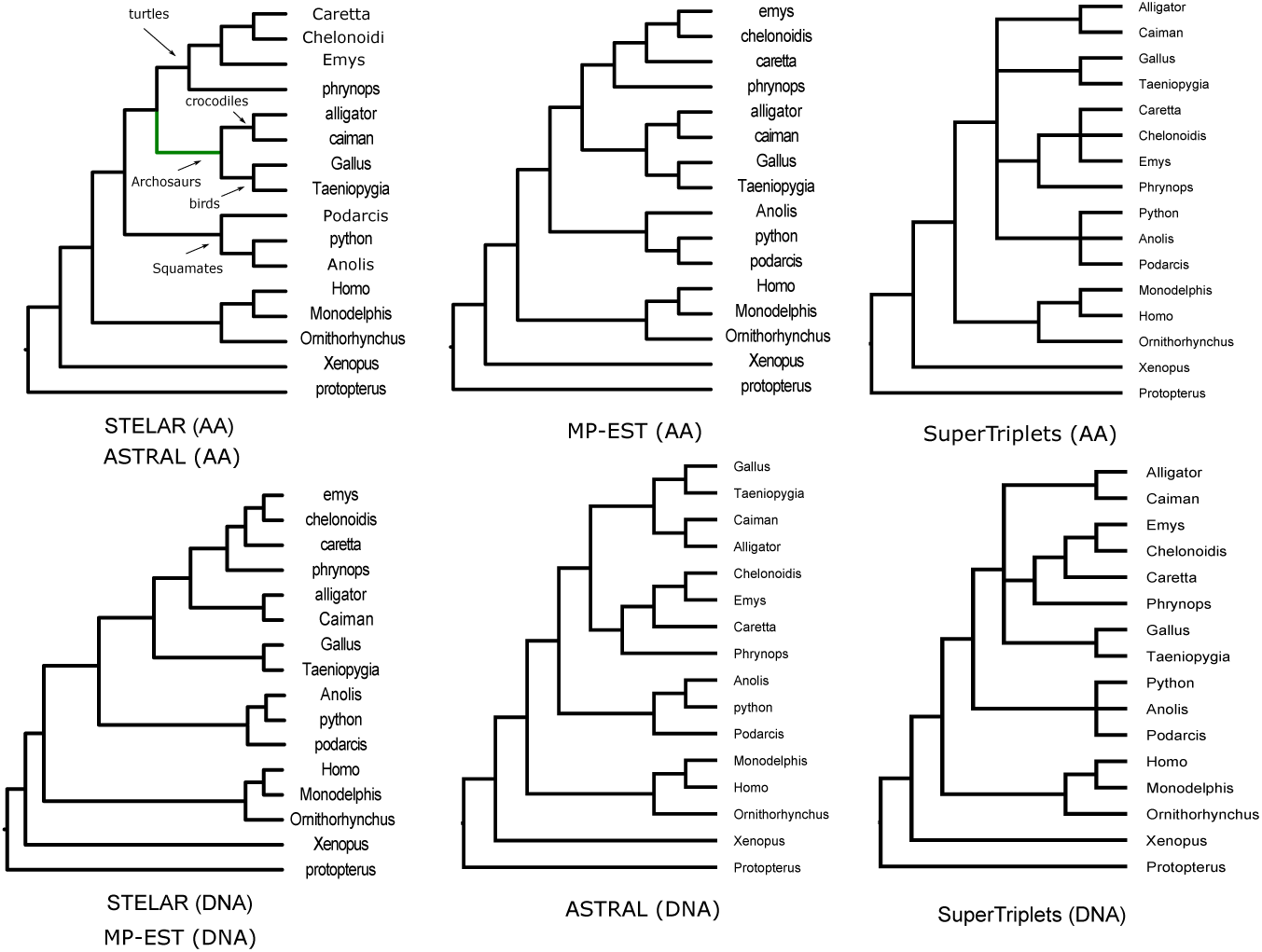
Analyses of the Amniota dataset (both DNA and AA gene trees) using STELAR, ASTRAL, MP-EST and SuperTriplets. We show the rooted version of the ASTRAL estimated trees using the outgroup (*Protopterus annectens*).

**Figure 6.**
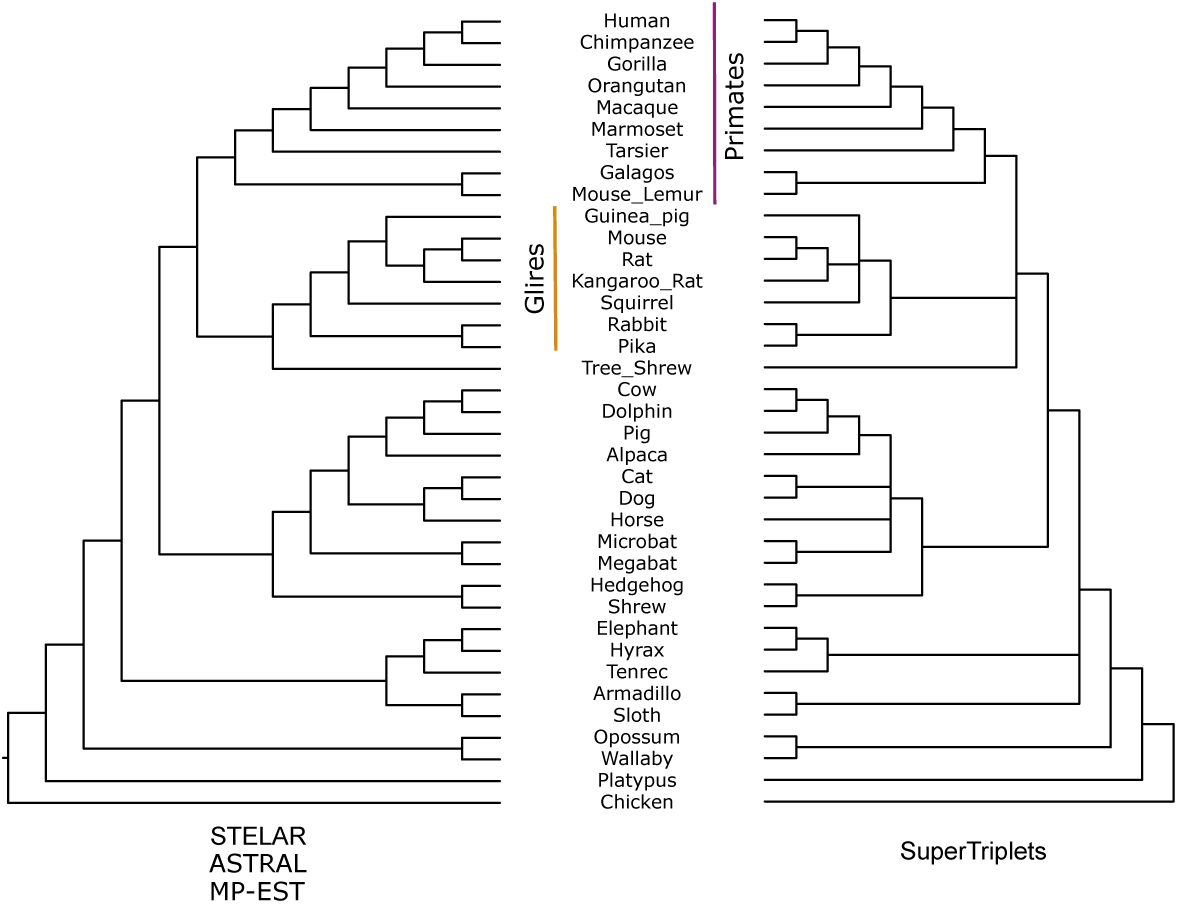
Analyses of the mammalian dataset using STELAR, ASTRAL, MP-EST and Super-Triplets.

### Running time

We ran the experiments on a Linux machine with 3.7 GiB RAM and 4 processors. Our experiments show that SuperTriplets is significantly faster than the other methods, and MP-EST is computationally more demanding than the others. For all the dataset analyzed in this study, SuperTriplets took a fraction of a second to run on a single replicate. STELAR and ASTRAL are substantially faster than MP-EST. The running time of MP-EST is more sensitive to the number of species than the number of genes, and grows rapidly as we increase the number of taxa [29]. On 37-taxon simulated dataset with 200 genes (moderate ILS), SuperTriplets took around half a second, both ASTRAL and STELAR took around 4 seconds whereas MP-EST (with 10 random starting points) took 3300 seconds (55 minutes). Running time of all these methods (except for SuperTriplets) increased with increasing amounts of ILS. On 37-taxon, 0.5X dataset (high ILS) with 200 genes, ASTRAL took around 10 seconds, STELAR took 15 seconds and MP-EST took 4800 seconds (80 minutes). This is because the numbers of distinct quartets and triplets in the gene trees increase as we increase the amount of discordance among the gene trees. When we increased the numbers of genes from 200 to 800 in 1X model condition, the running time of ASTRAL increased from 4 seconds to 30 seconds, whereas SuperTriplets took only 0.8 second and STELAR took 17 seconds. MP-EST took about 55 minutes and thus was not affected by the increased number of genes. On 11-taxon dataset, we ran the exact versions of ASTRAL and STELAR, and yet the running time is much less than MP-EST. Exact versions of ASTRAL and STELAR finished in less than a second on 5 *∼* 50 genes and less than two seconds on 50 *∼* 100 genes, whereas the running times of MP-EST range from 13 to 20 seconds. SuperTriplets took only 0.1 second irrespective of the numbers of genes. For 15-taxon dataset, MP-EST took 90 *∼* 100 seconds. ASTRAL and STELAR finished in less than 2 seconds on 100 gene dataset, and for 1000 genes they took about 50 and 80 seconds, respectively. The running time of SuperTriplets did not vary much with the change in the number of genes and finished in only around 0.4 second.

## Discussion and Conclusions

In this study, we presented STELAR – a method for estimating species trees from a set of estimated gene trees which is statistically consistent under the multi-species coalescent model. We formalized the constrained triplet consensus (CTC) problem and showed that the solution to the CTC problem is a statistically consistent estimator of the species tree under the MSC model. STELAR is a dynamic programming solution to the CTC problem. STELAR has an exact version (which provides the exact solution and takes exponential amount of time) as well as an efficient polynomial time heuristic version.

Extensive experimental studies on simulated dataset with varying model conditions suggest that STELAR, which is an efficient DP-based algorithm for solving the triplet consistency problem, can produce substantially better trees than SuperTriplets (which is a triplet-based supertre method and tries to find a median supertree with respect to a triplet dissimilarity measure), and MP-EST (which tries to maximize a pseudo-likelihood measure based on the triplet distribution of the gene trees). Thus, within the scope of the experiments conducted in this study, maximum triplet agreement has been shown to be a better optimization criterion than the triplet frequency based maximum pseudo-likelihood used in MP-EST, and the triplet dissimilarity based maximum parsimony criterion used in SuperTriplets, and as good as the maximum quartet agreement criterion used in ASTRAL. Experiments on Amniota and mammalian biological dataset suggest that STELAR can produce reliable trees on real biological dataset. Thus, we believe this study represents significant contributions, and STELAR would be considered as a useful tool in phylogenomics.

STELAR can consistently estimate the species tree topology and is fast enough to handle large numbers of genes and species, making it suitable for large scale phylogenomic analyses. However, STELAR can be expanded in several directions. Future work will need to investigate how the estimation of branch lengths (in coalescent unit) can be incorporated in STELAR. STELAR, at this stage, cannot handle unrooted and non-binary gene trees. The CTC problem for unrooted gene trees can be formulated as follows. The input is a set 𝓖= {*gt*_1_, *gt*_2_, *gt*_3_, …, *gt*_*k*_} of *k* unrooted gene trees, and the output is a species tree *ST* that maximizes the triplet consistency score with respect to a set 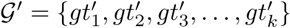 where 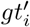 is a rooted version of *gt*_*i*_. That means, the idea is to find the optimal rooted versions of the unrooted gene trees so that the triplet agreement is maximized. Similar approach was previously applied [41] for estimating species trees from a set of unrooted gene trees by minimizing “extra lineages” resulting from deep coalescence. Similarly, the problem can be extended for non-binary gene trees as well. Thus, future works on formalizing the CTC problem for unrooted and non-binary gene trees and developing appropriate algorithms will be of great importance.

## Data and material availability

STELAR is freely available as an open source software at https://islamazhar.github.io/STELAR/. All the dataset analyzed in this paper are from previously published studies and are publicly available.

## Author’s contributions

MSB and RR conceived and designed the study; MI, KS and TD implemented STELAR; KS, TD, MI, MSB and RR proved the theoretical results; MI, KS and TD performed the experiments; MSB, KS, TD and MI wrote the first draft and all the authors took part in finalizing the manuscript.

